# A multi-continental analysis of the responses of freshwater food webs to climate and land use change

**DOI:** 10.1101/2022.04.18.488700

**Authors:** Gedimar Pereira Barbosa, Tadeu Siqueira

## Abstract

1. Food web responses to environmental change are not straightforward to understand as they occur through an intricate arrangement of direct and indirect effects. Although previous investigations have advanced knowledge on freshwater food web structure, we must better understand the intricate relationships between the main drivers of environmental change and trophic networks in lentic and lotic ecosystems.
2. We compiled multicontinental data to investigate how climate and land use change are related to the structure of freshwater food webs, considering the inherent differences in lentic and lotic ecosystems. We analyzed the direct and indirect relationships between land use intensity, and temperature and precipitation temporal trends, and food webs using multi-group structural equation modeling.
3. The strength and direction of the relationships between climate, land use, and food webs varied considerably among lentic and lotic ecosystems, but most indicated indirect effects through the number of links in the network. While network connectance both increased and decreased with land use and climate change, the number of trophic levels decreased with land use intensity and maximum temperature and increased with increasing precipitation. Omnivory increased with land use intensity in both ecosystems but was negatively related to changes on maximum temperature in lake food webs.
4. Even though food webs are expected to become more connected in face of disturbances, and our work supported this regarding local warming, the negative relationships between network connectance and land use intensification suggests that food webs become more specialized at disturbed sites. On the other hand, reduction in the number of trophic levels indicates the loss of top consumers in face of warming and increasing land use intensity.
5. The response of food webs in both lentic and lotic ecosystems to climate and land use change occurred mostly through changes in species interactions. Our results indicate that the intensification of land use makes food webs more specialized, with less trophic levels. Also, inherent aspects of freshwater ecosystems seemed to play a major role in the way food webs respond to disturbance and must be considered to fully understand and predict the effects of global changes on freshwater biodiversity.

## 1. INTRODUCTION

Food webs are rapidly transformed by environmental change (Estes et al., 2011; Tylianakis & Morris, 2017; Valiente-Banuet et al., 2015). While evidence shows that land use change disrupts food webs along a large range of ecosystems (Chen et al., 2021 2021; de Araújo et al., 2015; Jackson et al., 2020; Newbold et al., 2020), model predictions suggest that climate change might transform food webs to simpler structures with unprecedented consequences to ecosystem functioning and stability (Bartley et al., 2019; Lu et al., 2016; Thompson & Gonzalez, 2017). However, food web responses to environmental change are not always straightforward to understand as they occur through an intricate network of indirect and direct effects (Gibert, 2019). In addition, inherent aspects of ecosystems influence the response of food webs to environmental change, as already evidenced in lakes and rivers (Rosset et al., 2017). Even though previous investigations on the drivers of food web dynamics have made initial advances (Chen et al., 2021; Jackson et al., 2020; Thompson & Townsend, 2003), we must unravel the intricate network of factors that drive the response of lentic and lotic food webs to environmental change. Understanding how food webs relates to major global threats, such as land use intensification and changes in temperature and precipitation, could help identify controls of food web stability in lake and riverine ecosystems.

The conversion of natural to agricultural and urban landscapes affects food webs through changes in species composition and interactions (Tylianakis & Morris, 2017). As ecosystems are simplified and regions are homogenized, species composition changes due to the extinction of more sensitive species and the dominance of generalists that can overcome habitat alteration (Estes et al., 2011; Newbold et al., 2020). At the same time, more homogeneous ecosystems are also associated with higher interaction rates and the likelihood of finding prey, improving foraging success of generalist consumers (Laliberté & Tylianakis, 2010; Tylianakis et al., 2007). These changes may lead to more connected food webs because consumers that were once specialized start to coexist with all prey within their feeding niche (Riede et al., 2010). Food web structure also changes in face of cascading effects caused by agricultural and urban practices. For instance, while algae and invertebrates are directly influenced by agricultural pollution (Dubey & Dutta, 2020; Hayasaka et al., 2012; Rumschlag et al., 2020), species at higher trophic levels experience bottom-up effects, which may contribute to shortening food chain length due to the exclusion of more sensitive predators (Jackson et al., 2020; Newbold et al., 2020; Rumschlag et al., 2020). On the other hand, predator exclusion may lead to top-down effects, such as enhanced herbivory (Estes et al., 2011), and increased omnivory within the system, as omnivores have a broad foraging base and can survive where specialist predators cannot (Newbold et al., 2020; Wootton, 2017).

Land use change is also expected to influence food webs through interactions with changes in climate variables, especially temperature and precipitation (Jackson et al., 2020; Rosenblatt et al., 2017). Metabolic constraints imposed by warming increase the demand for food resources, leveling the rates of herbivory and predator-prey interactions (Hunt et al., 2017; O’Connor, 2009). With increased consumption and depletion of resources, warming can lead to the extinction of top consumers as large carnivores struggle to keep their metabolic demands and are replaced by omnivores or species at lower trophic levels (Jackson et al., 2020; Rosenblatt et al., 2017; Wootton, 2017). Warming may also unbalance interactions across trophic levels, causing contraction in the center of the trophic pyramid due to the predominance of tolerant and less palatable basal resources (Nagelkerken et al., 2020). However, the direct effects of warming on the structure of food webs may also depend on the predator’s ability to find and handle prey, in which more time spent searching and less time handling usually indicates consumer generalism, and thus, increased food web connectance (Petchey et al., 2010).

Food webs are strongly influenced by changes in precipitation amounts and regimes, because water limits the flux of biomass across trophic levels and govern the loss of species and interactions, particularly among predators (Ledger et al., 2013; Rosenblatt et al., 2017). Furthermore, the influence of precipitation and water availability on food webs also depend on physical aspects of different ecosystems (Rosset et al., 2017). For instance, in river catchments, altered precipitation is expected to influence food webs due to changes in flow regimes, an important aspect regulating primary productivity and ecosystem respiration (Bernhardt et al., 2022). On the other hand, under scenarios of reduced precipitation and increased drying conditions, lakes might suffer from reduced hydrological connectivity, which in turn influences food web structure due to changes on species diversity (Rosset et al., 2017). Lastly, altered precipitation and water deficits interact strongly with warming, amplifying the loss of biomass, reducing growth potential in species from all trophic levels, and increasing extinction rate of consumers at the top of food chains (Rosenblatt et al., 2017).

Despite the importance of water as the most essential resource for life, freshwater ecosystems are among the most affected ecosystems by land use intensification and climate change (Reid et al., 2019; Vörösmarty et al., 2010). Human population growth and increasing food demand have enclosed water bodies in agricultural and urban landscapes, affecting biodiversity through landscape simplification, depletion of natural resources and pollution (Foley, 2005; Vörösmarty et al., 2010). Also, as lentic and lotic ecosystems differ largely regarding to their physical, geochemical and biological aspects, it is essential to consider inherent ecosystem conditions on how freshwater communities respond to environmental disturbances (Rosset et al., 2017). Although quantifying the effects of land use on freshwater food webs in different parts of the globe is still challenging, advances in standardized databases on biological interactions (Brose et al., 2019; Poisot et al., 2016a), and large geospatial datasets (ESA, 2017; Abatzoglou et al., 2018) offer reliable ways to the investigation of the effects of environmental change on species and their interactions in lakes, ponds, rivers, and streams.

Here, we carefully compiled data on 51 freshwater food webs sampled in 12 countries and six continents to investigate how climate and land use change are related to the structure of freshwater food webs, considering the inherent differences in lentic and lotic ecosystems. For this, we analyzed the direct and indirect relationships between land use intensity, and temperature and precipitation temporal trends, and food webs using multi-group structural equation modeling (multi-group SEM) (Supporting Fig. S1). As land use intensification leads to more homogenous and less structured ecosystems, we expected that the number of nodes would decrease and the number of links increase with land use intensification, leading to more connected food webs (Laliberté & Tylianakis, 2010; Riede et al., 2010; Tylianakis et al., 2007). We also expected food webs under intense land use and warming temperatures to have lower trophic levels due to the extinction of top predators (Estes et al., 2011), increased control of basal resources by enhanced herbivory (O’Connor, 2009), and higher omnivory (Wootton, 2017). However, as omnivores can shift their diet and replace lost specialized carnivores, occupying newly available trophic levels (Newbold et al., 2020), overall presence of omnivory within food webs could also be reduced under environmental changes. Because water availability plays an important role in food web structure (Jackson et al., 2020; Rosenblatt et al., 2017), we expected precipitation to enhance the ability of food webs in reaching higher trophic levels. Lastly, we also predicted land use intensity to drive food web structure through its effects on temperature and precipitation.

## 2. METHODS

### 2.1. Food web Data

We obtained freshwater food webs from three sources: the Mangal interaction database, using the *rmangal* package in R (Poisot et al., 2016a), the GlobAl databasE of traits and food Web Architecture (GATEWAY) version 1.0 (Brose et al., 2019), and the Interaction Web Data Base (IWDB) (http://www.ecologia.ib.usp.br/iwdb/index.html). As lake food webs meeting our criteria were rarely found in these databases, we also performed a search in the Web of Science Core Collection (see Supporting Information for searched terms). Food webs usually vary in sampling procedures, particularly among lentic and lotic ecosystems. Thus, we kept only those in which researchers reported the occurrence of predation, herbivory, and detritivory interactions. In cases where sites were sampled more than once, resulting in replicated data for the same location, we selected those with higher resolution (number of nodes). We checked the original publications for geographical location and sampling date when information was not available in the databases. Because the land use database we used contained land cover data from 1992 to 2015, we only selected food webs sampled within this period. We considered the sampling date as the year in which the researchers concluded field sampling. To calculate food web metrics, we transformed original data into adjacent matrices (*A* = [*i, j*]), in which *i* represents the resources and *j* represents consumers.

### 2.2. Environmental Variables

To test the relationship between food web properties and land use intensity, we first characterized land use on the surroundings of the sites where food webs were originally sampled. We did that by extracting land cover information from the global ESA CCI database (ESA, 2017), an annually generated land cover product at 300 m resolution for the period 1992 – 2015. The data set comprises 38 land cover categories, following the classification system developed by the United States Food and Agricultural Organization (ESA, 2017). To extract cell values, we established a 1 km buffer around the locations where food webs were sampled. In cases of food webs sampled in large water bodies, such as lakes, we extended the 1 km buffer to the total surrounding area of the water body (see Fig. S2 in the Supporting Information for an example). To estimate land use intensity, we attributed a value of 1.0 to cells falling into urban areas (category 190 in the ESA CCI database) and cropland areas (categories 10, 11, 12, 30, and 40), and 0.0 to cells pertaining to other categories (e.g., forest and grassland areas). Because categories 30 and 40 are mixed cropland and natural vegetation (30 – mosaic (>50%) cropland and (<50%) natural vegetation, and 40 -mosaic (>50%) natural vegetation and (<50%) cropland), their cell values were down-weighted to 0.75 and 0.25, respectively. The values reported in the analysis represent the average of land use within the 1 km buffer.

To quantify climate change, we gathered temperature and precipitation data from the TerraClimate database, a monthly generated product for climate and climatic water balance for global terrestrial surfaces at ∼ 4 km for the period 1958 – 2015 (Abatzoglou et al., 2018). We extracted monthly mean values of maximum temperature and precipitation at the exact cell coordinates where food webs were sampled, following a timeframe of thirty years past to the year food webs were sampled. Then, we estimated temporal trends for each variable using generalized additive models, accounting for a temporally autocorrelated structure with the R package *mgcv* (Wood, 2017). Finally, we extracted the linear slope from the model for each variable at each site, which summarized the trend for maximum temperature and precipitation. We selected maximum temperature and precipitation because these variables are better representations of the warming and water deficit conditions considered in our hypothesis.

### 2.3. Food Web Variables

Although a variety of metrics to describe the structure of food webs are available (Delmas et al., 2019), here we measured only aspects related to our hypothesis. First, we quantified the basic properties of food webs, as number of nodes: ranging from species level to broader resolution groups (e.g., algae, detritus), number of links: the realized interactions between nodes in the food web, and proportion of basal species: the fraction of nodes that represent primary producers. Then, we calculated network level responses, such as mean omnivory: the average value of the omnivory index calculated for each node within the food web, maximum trophic level: the highest level obtained for the nodes within the food web, and connectance: calculated here, as the residuals from a linear regression between the number of realized interactions and the number of possible ones (both log-transformed). Omnivory and trophic level indexes were calculated using the package *NetIndices* (Kones et al., 2009).

### 2.4. Data Analysis

Because land use intensification might influence food web structure through distinct pathways, we used piecewise structural equation modeling (SEM) to test our hypothesis (Lefcheck, 2016). For this, we used ecological theory and knowledge of the system to define the paths (Supporting Fig. S1), and fit linear mixed-effects models to decompose the relative contribution of explanatory variables to the structural responses of food webs to land use and climate, both direct and indirect (see the Supporting Information for model description). The term “direct” here means a relationship not mediated by other variables in the SEM, but that can rely on mechanisms or variables not included in this work. To account for variation among food webs caused by differences in methodological procedures (Thompson et al., 2001), we incorporated sampling method (modeling, gut-content, and gut-isotope analysis) as a random factor within the mixed models. Overall model fit was evaluated using the Shipley d-separation test, which produces a Fisher’s *C* statistic to a chi-square distribution (Shipley, 2000). To directly compare paths between lentic and lotic ecosystems, we employed a multi-group analysis to our SEM. The multi-group piecewise SEM indicates which paths change based on groups, allowing direct comparison between groups within the same model. To ensure that differences in the number of lentic and lotic food webs compiled would not affect our model, we performed a sensitivity analysis in which we used the same number of lentic and lotic food webs by fitting the model on 999 networks randomly selected from our dataset (see the Supporting Information for details). We conducted the piecewise SEM using the R package *piecewiseSEM* (Lefcheck, 2016), and for the linear mixed models, we used the function *lme* in the R package *nlme* (Pinheiro et al., 2021).

## 3. RESULTS

Our search resulted in 51 freshwater food webs, averaging 59 nodes and 196 links, sampled in 12 countries and six continents (Fig. 1, Table S1 in Supporting Information). As expected, rivers and streams accounted for most of the food webs in our dataset (n = 37, against 14 in lakes and ponds). However, differences in the number of lentic and lotic food webs did not affect model results, as demonstrated by our sensitivity analysis (Fig. S3 in Supporting Information). Overall, food web structure varied considerably among ecosystems (Fig. 2), especially regarding to the number of nodes and proportion of basal species, which showed almost an asymmetry between lentic and lotic food webs. By contrast, the number of realized interactions followed a similar pattern in both ecosystem types, with most food webs peaking around 100 and 200 links for lentic and lotic ecosystems, respectively. This discrepancy in the distribution of nodes and links between ecosystems was also evidenced by the distribution of connectance, in which lotic food webs showed a more uniform distribution in comparison to lentic ones. Most food webs in rivers and streams reached around three trophic levels and showed a low number of omnivores, when compared to a more homogeneous distribution of omnivory and maximum trophic levels found in lakes and ponds.

**Figure 1.**
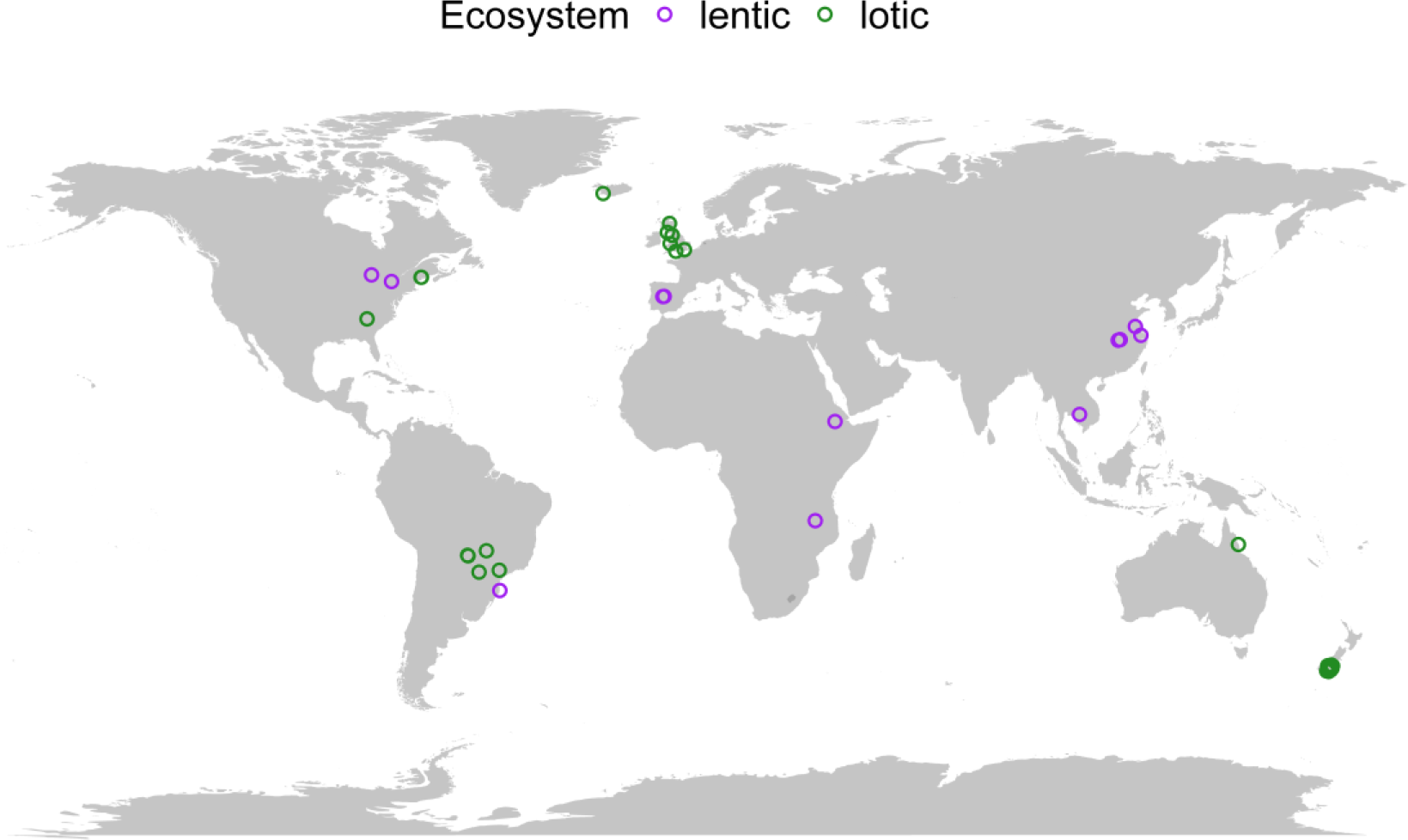
Geographical distribution of the freshwater food webs used in this study. Green and purple circles indicate lotic and lentic food webs, respectively. Coordinates are based on data set meta-data and original publications. Central points are considered for large water bodies (e.g., Lake Ontario). Sites in China, New Zealand and Spain are indistinguishable at this map scale.

**Figure 2.**
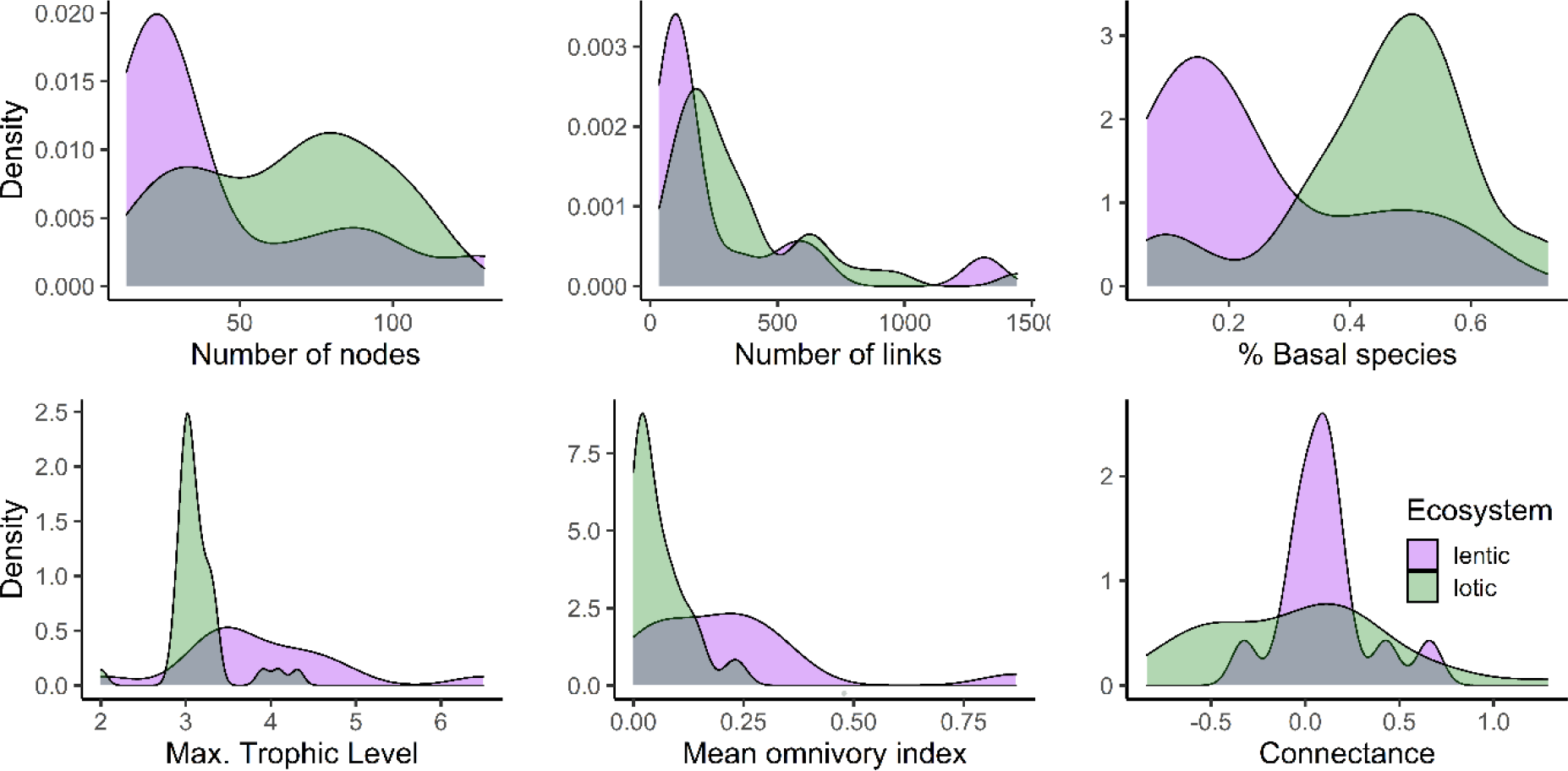
Adjusted density estimates of food web metrics calculated for both ecosystems in our multi-continental data set. Green and purple areas indicate lotic and lentic food webs, respectively.

Overall, multi-group SEM fit the data very well (*C*14 = 11.548, *P* = 0.643, Fig. 3), and revealed contrasting associations of freshwater food webs to environmental predictors. Even though some food web variables were directly linked to land use intensity and climate change, as expected, most relationships between environmental predictors and food webs occurred through indirect paths mediated by the number of nodes and links (Table 1). The model also showed how the relationship between freshwater food webs and environmental changes is mediated by ecosystem type, particularly the relationship between land use intensity and food web structure, and precipitation and food web structure. However, ecosystem differences did not influence the associations between maximum temperature and food web variables. In total, nine relationships were not constrained to the global model (Fig. 3).

**Figure 3.**
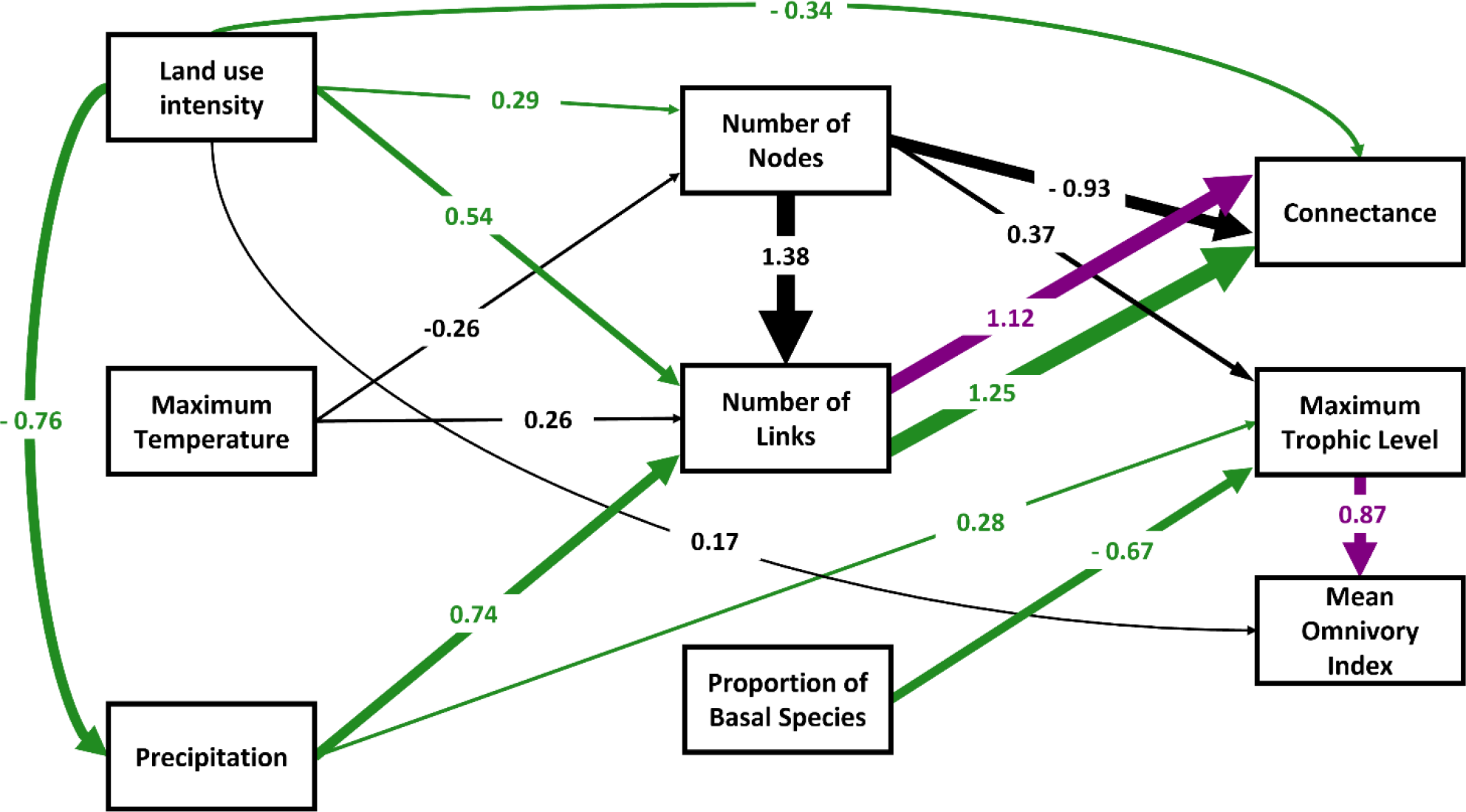
Multi-group SEM that relates land use intensity and climate variables to the structure of freshwater food webs across continents. Green, purple and black arrows represent significant relationships between endogenous and exogenous variables in lotic, lentic and in both ecosystems, respectively. Arrow width represents the magnitude of relationship, following standardized coefficients (*β*) reported along the connections. Non-significant relationships were hidden for clarity (see Supporting Fig. S1 for full model).

**Table 1.** Direct and indirect relationships between land use intensity and climate change and food web structure in lotic and lentic ecosystems. For a detailed description of how coefficients were calculated, see Supporting Information.

| Ecosystem | Predictors | Food web structure | Nº Paths | Direct coef. | Indirect coef. | Total coef. |
| --- | --- | --- | --- | --- | --- | --- |
| Lotic | Land use intensity | Connectance | 5 | -0.34 | 0.20 | -0.14 |
| Lotic | Land use intensity | Max. Trophic level | 2 | 0.00 | -0.11 | -0.11 |
| Lotic | Land use intensity | Mean omnivory index | 1 | 0.17 | 0.00 | 0.17 |
| Lotic | Max. Temperature | Connectance | 3 | 0.00 | 0.12 | 0.12 |
| Lotic | Max. Temperature | Max. Trophic level | 1 | 0.00 | -0.10 | -0.10 |
| Lotic | Precipitation | Connectance | 1 | 0.00 | 0.93 | 0.93 |
| Lotic | Precipitation | Max. Trophic level | 1 | 0.28 | 0.00 | 0.28 |
| Lentic | Land use intensity | Mean omnivory index | 2 | 0.17 | 0.00 | 0.17 |
| Lentic | Max. Temperature | Connectance | 3 | 0.00 | 0.13 | 0.13 |
| Lentic | Max. Temperature | Max. Trophic level | 1 | 0.00 | -0.10 | -0.10 |
| Lentic | Max. Temperature | Mean omnivory index | 1 | 0.00 | -0.08 | -0.08 |

Although there was not a relationship between land use intensity and connectance in lakes or ponds, connectance in lotic ecosystems was negatively related to land use intensity through direct and indirect paths (Fig. 3; Table 1). When we considered only an indirect relationship through the number of links, land use intensity was positively related to food web connectance, as we predicted. This was also the case for the relationship between connectance and climate variables, with indirect paths showing that connectance was both positively and negatively related to maximum temperature, depending on the indirect path (Fig. 3). Despite that, the resulting relationship between maximum temperature and connectance was negative for both ecosystems (Table 1). Precipitation, in contrast, was only related to lotic food webs properties (Fig.3), and, against our expectations, was positively related to connectance through the number of links (Table 1).

Our model also showed divergent relationships between land use intensity and maximum trophic level, with a positive relationship in both ecosystems mediated by the number of nodes, and a negative relationship mediated by precipitation only in lotic food webs (Fig. 3; Table 1). As we predicted, maximum trophic level was negatively related to maximum temperature through an indirect relationship with the number of nodes, in both lentic and lotic ecosystems, and positively related to increased precipitation only in rivers and streams (Fig. 3). While omnivory increased with land use intensity in both ecosystems, it slightly decreased with maximum temperature only in lentic food web (Fig. 3; Table 1).

The environmental predictors explained a large portion of the variation in some food web metrics (Number of Links = 0.69; Connectance = 0.77; Mean Omnivory Index = 0.84; Table 2). As shown by marginal and conditional *R2* values, food web inference methods (modeling, gut-content, and gut-isotope analysis), which were included in the SEM as random effects, did not affect the relationships between predictors and food web metrics, such as connectance, maximum trophic level and mean omnivory index. On the other hand, the amount of variation explained in % of basal species and number of nodes was almost entirely due to the sampling method used to infer trophic interactions (Table 2).

**Table 2.** Marginal and Conditional *R*^*2*^ values obtained for individual endogenous variables, following the multi-group SEM that relates freshwater food webs to land use intensity and climate change.

| Endogenous Variables | Marginal $R^2$ | Conditional $R^2$ |
| --- | --- | --- |
| Max. Temperature | 0.00 | 0.36 |
| % of Basal Species | 0.01 | 0.76 |
| Number of Nodes | 0.08 | 0.69 |
| Precipitation | 0.14 | 0.20 |
| Max. Trophic Level | 0.21 | 0.64 |
| Number of Links | 0.69 | 0.89 |
| Connectance | 0.77 | 0.77 |
| Mean Omnivory Index | 0.84 | 0.86 |

## 4. DISCUSSION

In general, our models indicate that freshwater food webs were strongly influenced by changes in temperature and precipitation that occurred in the last 30 years and by land use intensity within the river basin. However, the relationships we observed were sometimes conflicting and relied deeply on our ability to detect indirect paths involving environmental changes and food web structure. Also, as lentic and lotic ecosystems differ fundamentally in their structure, dynamics and in the way they are affected by the surrounding landscape, food webs responded differently to climate and land use changes. More specifically, our models indicate that food web structure in streams and rivers were more strongly related to land use intensity and climate variables, than those in lakes and ponds.

We predicted that food web connectance would increase with land use intensity, based on the assumption that as ecosystem complexity is reduced in disturbed landscapes (Estes et al., 2011), generalists replace specialists, thus increasing the number of realized interactions (Laliberté & Tylianakis, 2010; Riede et al., 2010; Tylianakis et al., 2007). This prediction was partially supported for lotic food webs, but only when considering indirect paths linking land use intensity and connectance through the number of links. However, the overall relationship, as indicated by the multiplied path coefficients, indicate that connectance decreases with land use intensity. This suggests that food webs become more specialized at the local scale, following the removal of specialists and more susceptible species, reduced competition, and restricted food choices (de Araújo et al., 2015).

The relationship between warming and connectance was consistent among lentic and lotic ecosystems, suggesting that freshwater food webs are becoming more connected as temperature rises. This agrees with previous findings (Gibert, 2019; Petchey et al., 2010) in which temperature was positively related to food web connectance. High temperatures impose metabolic constrains to species and level up consumption rates within food webs (O’Connor, 2009). Such scenario can lead to depletion of resources and, consequently, to the local extinction of functional feeding groups (Rosenblatt et al., 2017). Our results support these ideas, as warming was indirectly related to connectance through a negative relationship with the number of nodes, and a positive relationship with the number of links in the food web. Precipitation, on the other hand, was related to connectance only in rivers and streams, via an indirect path through number of links. This is somehow expected because, under scenarios of reduced precipitation, physiological stress induces consumer-specific shifts in diet due to modified foraging strategies or changes in the abundance and distribution of resources (Ledger et al., 2013; Lu et al., 2016). Although this finding shows that ecosystem differences play a role in the relationship between precipitation and food web structure, future work is needed to elucidate the mechanism behind it.

While temperature was the only predictor related to maximum trophic levels in lakes and ponds, temperature, precipitation, and land use intensity were all related to maximum trophic levels in lotic ecosystems. Land use intensity and increased temperature were related to food webs with lower trophic levels, which might be an indicative of shortening food chain length due to predator exclusion (Ledger et al., 2013; Newbold et al., 2020), as already found in stream food webs (Jackson et al., 2020). On the other hand, food webs under increased precipitation reached higher trophic levels, which is expected because water availability enhances biomass production within the system (Rosenblatt et al., 2017), thus, supporting higher food chain lengths (Ziegler et al., 2015). Our results also support the assumption of omnivory being more common in disturbed ecosystems (Wootton, 2017), as land use intensity was positively related to mean omnivory index, despite ecosystem differences. In lakes and ponds, however, the negative influence of temperature on maximum trophic level drove a slightly negative relationship between warming and omnivory. This indirect relationship provides some support to the idea of omnivores replacing specialized consumers, when facing disturbances caused by climate changes.

Although we carefully selected the data included in this study, some caveats need to be considered. For example, our analyses included limited data from Africa, Asia, and some parts of Europe. Thus, although our analysis includes information from 6 continents, our inferences should not be seen as global. Also, food web sampling methods differ largely among primary studies, and even though we reduced some of the variability by including sampling method as random effects in our model, potential biases arise from variables we could not control, such as sampling period and taxonomic resolution. Despite this, considering the number of data sets we included in our analysis and the strength of the relationships we found, we consider that our results help advancing our understanding of the effects of environmental changes on the structure of freshwater food webs.

In summary, our analyses reveal that fundamental aspects of food web structure, such as connectance and the number of trophic levels, vary considerably with climate and land use change. This is alarming, because the conversion of natural to modified landscapes has been happening rapidly (Ellis et al., 2021). We also found that the number of links in a food web plays a major role as a mediator linking complex relationships between environmental drivers and food web structure. This reinforces the idea that interactions are oftentimes decoupled from species richness, and thus can be lost before species become locally extinct, risking the maintenance of essential ecological functions (Valiente-Banuet et al., 2015). Such decoupled dynamics between interactions and species loss also strengthens the importance of considering ecological networks to understand and predict biodiversity changes (Poisot et al., 2016b). Lastly, as our results implicitly show, inherent aspects of freshwater ecosystems play a major role in the way food webs are structured and respond to disturbance and, thus, must be considered to fully understand the effects of environmental change on freshwater biodiversity.

## ACKNOWLEDGEMENTS

This study was financed in part by the Coordenação de Aperfeiçoamento de Pessoal de Nível Superior-Brasil (CAPES) - Finance Code 001 through a PhD scholarship conceived to GPB. TS was supported by grant #309496/2021-7, Conselho Nacional de Desenvolvimento Científico e Tecnológico (CNPq) and grant 21/00619-7, São Paulo Research Foundation (FAPESP). We thank Mathias Pires and Míriam Pilz Albrecht for providing valuable comments on an earlier version of this manuscript.

## DATA AVAILABILITY STATEMENT

All data and code used to reproduce the analysis are available in Zenodo (https://doi.org/10.5281/zenodo.6501835)

## REFERENCES

Abatzoglou, J. T., Dobrowski, S. Z., Parks, S. A., & Hegewisch, K. C. (2018). TerraClimate, a high-resolution global dataset of monthly climate and climatic water balance from 1958–2015. Scientific Data, 5(1), 170191. https://doi.org/10.1038/sdata.2017.191

Bartley, T. J., McCann, K. S., Bieg, C., Cazelles, K., Granados, M., Guzzo, M. M., … McMeans, B. C. (2019). Food web rewiring in a changing world. Nature Ecology & Evolution, 3(3), 345–354. https://doi.org/10.1038/s41559-018-0772-3

Bernhardt, E. S., Savoy, P., Vlah, M. J., Appling, A. P., Koenig, L. E., Hall, R. O., … Grimm, N. B. (2022). Light and flow regimes regulate the metabolism of rivers. Proceedings of the National Academy of Sciences, 119(8), e2121976119. https://doi.org/10.1073/pnas.2121976119

Brose, U., Archambault, P., Barnes, A. D., Bersier, L.-F., Boy, T., Canning-Clode, J., … Iles, A. C. (2019). Predator traits determine food-web architecture across ecosystems. Nature Ecology & Evolution, 3(6), 919–927. https://doi.org/10.1038/s41559-019-0899-x

Chen, Z., Dudgeon, D., & Liew, J. H. (2021). Human settlements in headwater catchments are associated with generalist stream food webs. Hydrobiologia, 848(17), 4017– 4027. https://doi.org/10.1007/s10750-021-04620-y

de Araújo, W. S., Vieira, M. C., Lewinsohn, T. M., & Almeida-Neto, M. (2015). Contrasting Effects of Land Use Intensity and Exotic Host Plants on the Specialization of Interactions in Plant-Herbivore Networks. PLOS ONE, 10(1), e0115606. https://doi.org/10.1371/journal.pone.0115606

Delmas, E., Besson, M., Brice, M.-H., Burkle, L. A., Dalla Riva, G. V., Fortin, M.-J., … Poisot, T. (2019). Analysing ecological networks of species interactions: Analyzing ecological networks. Biological Reviews, 94(1), 16–36. https://doi.org/10.1111/brv.12433

Dubey, D., & Dutta, V. (2020). Nutrient Enrichment in Lake Ecosystem and Its Effects on Algae and Macrophytes. In V. Shukla & N. Kumar (Eds.), Environmental Concerns and Sustainable Development (pp. 81–126). Singapore: Springer Singapore. https://doi.org/10.1007/978-981-13-6358-0_5

Ellis, E. C., Gauthier, N., Klein Goldewijk, K., Bliege Bird, R., Boivin, N., Díaz, S., … Watson, J. E. M. (2021). People have shaped most of terrestrial nature for at least 12,000 years. Proceedings of the National Academy of Sciences, 118(17), e2023483118. https://doi.org/10.1073/pnas.2023483118

ESA (2017). Land Cover CCI Product User Guide Version 2. Tech. Rep. maps.elie.ucl.ac.be/CCI/viewer/download/ESACCI-LC-Ph2-PUGv2_2.0.pdf

Estes, J. A., Terborgh, J., Brashares, J. S., Power, M. E., Berger, J., Bond, W. J., … Wardle, D. A. (2011). Trophic Downgrading of Planet Earth. Science, 333(6040), 301–306. https://doi.org/10.1126/science.1205106

Foley, J. A. (2005). Global Consequences of Land Use. Science, 309(5734), 570–574. https://doi.org/10.1126/science.1111772

Gibert, J. P. (2019). Temperature directly and indirectly influences food web structure. Scientific Reports, 9(1), 5312. https://doi.org/10.1038/s41598-019-41783-0

Hayasaka, D., Korenaga, T., Suzuki, K., Saito, F., Sánchez-Bayo, F., & Goka, K. (2012). Cumulative ecological impacts of two successive annual treatments of imidacloprid and fipronil on aquatic communities of paddy mesocosms. Ecotoxicology and Environmental Safety, 80, 355–362. https://doi.org/10.1016/j.ecoenv.2012.04.004

Hunt, S. K., Galatowitsch, M. L., & McIntosh, A. R. (2017). Interactive effects of land use, temperature, and predators determine native and invasive mosquito distributions. Freshwater Biology, 62(9), 1564–1577. https://doi.org/10.1111/fwb.12967

Jackson, M. C., Fourie, H. E., Dalu, T., Woodford, D. J., Wasserman, R. J., Zengeya, T. A., … Weyl, O. L. F. (2020). Food web properties vary with climate and land use in South African streams. Functional Ecology, 34(8), 1653–1665. https://doi.org/10.1111/1365-2435.13601

Kones, J. K., Soetaert, K., van Oevelen, D., & Owino, J. O. (2009). Are network indices robust indicators of food web functioning? A Monte Carlo approach. Ecological Modelling, 220(3), 370–382. https://doi.org/10.1016/j.ecolmodel.2008.10.012

Laliberté, E., & Tylianakis, J. M. (2010). Deforestation homogenizes tropical parasitoid– host networks. Ecology, 91(6), 1740–1747. https://doi.org/10.1890/09-1328.1

Ledger, M. E., Brown, L. E., Edwards, F. K., Milner, A. M., & Woodward, G. (2013). Drought alters the structure and functioning of complex food webs. Nature Climate Change, 3(3), 223–227. https://doi.org/10.1038/nclimate1684

Lefcheck, J. S. (2016). PIECEWISESEM: Piecewise structural equation modelling in R for ecology, evolution, and systematics. Methods in Ecology and Evolution, 7(5), 573– 579. https://doi.org/10.1111/2041-210X.12512

Lu, X., Gray, C., Brown, L. E., Ledger, M. E., Milner, A. M., Mondragón, R. J., … Ma, A. (2016). Drought rewires the cores of food webs. Nature Climate Change, 6(9), 875– 878. https://doi.org/10.1038/nclimate3002

Nagelkerken, I., Goldenberg, S. U., Ferreira, C. M., Ullah, H., & Connell, S. D. (2020). Trophic pyramids reorganize when food web architecture fails to adjust to ocean change. Science, 369(6505), 829–832. https://doi.org/10.1126/science.aax0621

Newbold, T., Bentley, L. F., Hill, S. L. L., Edgar, M. J., Horton, M., Su, G., … Purvis, A. (2020). Global effects of land use on biodiversity differ among functional groups. Functional Ecology, 34(3), 684–693. https://doi.org/10.1111/1365-2435.13500

O’Connor, M. I. (2009). Warming strengthens an herbivore–plant interaction. Ecology, 90(2), 388–398. https://doi.org/10.1890/08-0034.1

Petchey, O. L., Brose, U., & Rall, B. C. (2010). Predicting the effects of temperature on food web connectance. Philosophical Transactions of the Royal Society B: Biological Sciences, 365(1549), 2081–2091. https://doi.org/10.1098/rstb.2010.0011

Pinheiro, J., Bates, D., DebRoy, S., Sarkar, D., & R Core Team (2021). nlme: Linear and Nonlinear Mixed Effects Models. R package version 3.1-152, https://CRAN.R-project.org/package=nlme.

Poisot, T., Baiser, B., Dunne, J. A., Kéfi, S., Massol, F., Mouquet, N., … Gravel, D. (2016a). mangal—Making ecological network analysis simple. Ecography, 39(4), 384–390. https://doi.org/10.1111/ecog.00976

Poisot, T., Stouffer, D. B., & Kéfi, S. (2016b). Describe, understand and predict: Why do we need networks in ecology? Functional Ecology, 30(12), 1878–1882. https://doi.org/10.1111/1365-2435.12799

Reid, A. J., Carlson, A. K., Creed, I. F., Eliason, E. J., Gell, P. A., Johnson, P. T. J., … Cooke, S. J. (2019). Emerging threats and persistent conservation challenges for freshwater biodiversity. Biological Reviews, 94(3), 849–873. https://doi.org/10.1111/brv.12480

Riede, J. O., Rall, B. C., Banasek-Richter, C., Navarrete, S. A., Wieters, E. A., Emmerson, M. C., … Brose, U. (2010). Scaling of Food-Web Properties with Diversity and Complexity Across Ecosystems. Chapter 3. In: Advances in Ecological Research (Vol. 42, pp. 139–170). Elsevier. https://doi.org/10.1016/B978-0-12-381363-3.00003-4

Rosenblatt, A. E., Smith-Ramesh, L. M., & Schmitz, O. J. (2017). Interactive effects of multiple climate change variables on food web dynamics: Modeling the effects of changing temperature, CO2, and water availability on a tri-trophic food web. Food Webs, 13, 98–108. https://doi.org/10.1016/j.fooweb.2016.10.002

Rosset, V., Ruhi, A., Bogan, M. T., & Datry, T. (2017). Do lentic and lotic communities respond similarly to drying? Ecosphere, 8(7), e01809. https://doi.org/10.1002/ecs2.1809

Rumschlag, S. L., Mahon, M. B., Hoverman, J. T., Raffel, T. R., Carrick, H. J., Hudson, P. J., & Rohr, J. R. (2020). Consistent effects of pesticides on community structure and ecosystem function in freshwater systems. Nature Communications, 11(1), 6333. https://doi.org/10.1038/s41467-020-20192-2

Shipley, B. (2000). A New Inferential Test for Path Models Based on Directed Acyclic Graphs. Structural Equation Modeling: A Multidisciplinary Journal, 7(2), 206–218. https://doi.org/10.1207/S15328007SEM0702_4

Thompson, P. L., & Gonzalez, A. (2017). Dispersal governs the reorganization of ecological networks under environmental change. Nature Ecology & Evolution, 1(6), 0162. https://doi.org/10.1038/s41559-017-0162

Thompson, R. M., Edwards, E. D., McIntosh, A. R., & Townsend, C. R. (2001). Allocation of effort in stream food-web studies: The best compromise? Marine and Freshwater Research, 52(3), 339. https://doi.org/10.1071/MF00041

Thompson, R. M., & Townsend, C. R. (2003). IMPACTS ON STREAM FOOD WEBS OF NATIVE AND EXOTIC FOREST: AN INTERCONTINENTAL COMPARISON. Ecology, 84(1), 145–161. https://doi.org/10.1890/0012-9658(2003)084[0145:IOSFWO]2.0.CO;2

Tylianakis, J. M., & Morris, R. J. (2017). Ecological Networks Across Environmental Gradients. Annual Review of Ecology, Evolution, and Systematics, 48(1), 25–48. https://doi.org/10.1146/annurev-ecolsys-110316-022821

Tylianakis, J. M., Tscharntke, T., & Lewis, O. T. (2007). Habitat modification alters the structure of tropical host–parasitoid food webs. Nature, 445(7124), 202–205. https://doi.org/10.1038/nature05429

Valiente-Banuet, A., Aizen, M. A., Alcántara, J. M., Arroyo, J., Cocucci, A., Galetti, M., … Zamora, R. (2015). Beyond species loss: The extinction of ecological interactions in a changing world. Functional Ecology, 29(3), 299–307. https://doi.org/10.1111/1365-2435.12356

Vörösmarty, C. J., McIntyre, P. B., Gessner, M. O., Dudgeon, D., Prusevich, A., Green, P., … Davies, P. M. (2010). Global threats to human water security and river biodiversity. Nature, 467(7315), 555–561. https://doi.org/10.1038/nature09440

Wood, S. N. (2017). Generalized Additive Models: An Introduction with R (2nd ed.). New York, NW: Chapman and Hall/CRC.

Wootton, K. L. (2017). Omnivory and stability in freshwater habitats: Does theory match reality? Freshwater Biology, 62(5), 821–832. https://doi.org/10.1111/fwb.12908

Ziegler, J. P., Solomon, C. T., Finney, B. P., & Gregory-Eaves, I. (2015). Macrophyte biomass predicts food chain length in shallow lakes. Ecosphere, 6(1), art5. https://doi.org/10.1890/ES14-00158.1

