## Supporting Information for "A multi-continental analysis of the responses of freshwater food webs to climate and land use change"

#### 1. Model Description and Analysis

We used a hypothetical model to evaluate the effects of land use intensity and climate change on the structure of food webs (Fig. S1). Food web sampling method (modeling, gut-content, and gut-isotope analysis) was considered as random effect within the model.

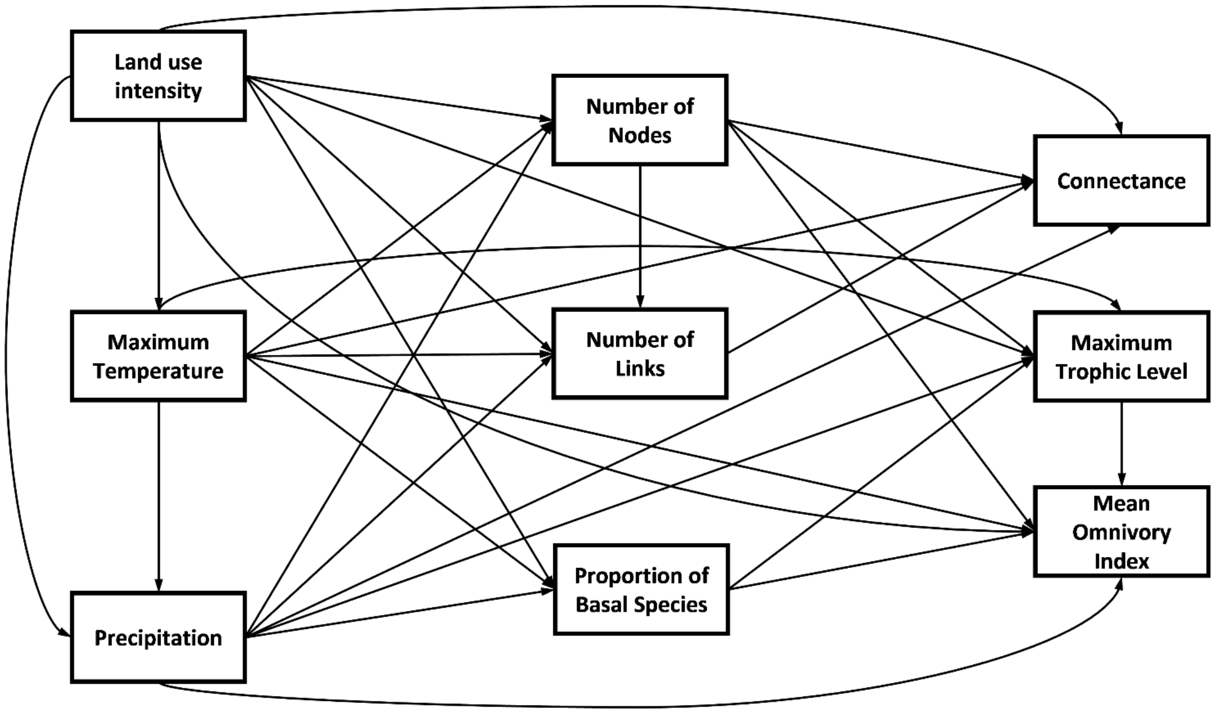


**Fig. S1.** Hypothetical model for the effects of land use intensity and climate change on the structure of freshwater food webs

A standard regression coefficient ($\beta$) was retrieved for each relationship between exogenous and endogenous variables, using *piecewiseSEM*. Then, indirect path coefficients were obtained as follows:

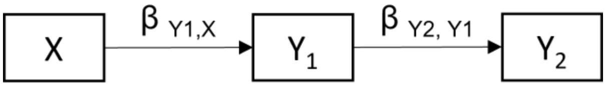

Considering $\beta_{Y1,X}$ as the standard regression coefficient between $X$ and $Y_1$, and $\beta_{Y2,Y1}$ as the standard regression coefficient between $Y_1$ and $Y_2$, the indirect effect of $X$ on $Y_2$ was obtained by:

$$\beta_{Y2,X} = \beta_{Y1,X} * \beta_{Y2,Y1}$$

After calculating the direct and indirect coefficients, we obtained the total effect of a predictor on a response variable by summing all effects from direct and indirect paths.

### 2. Food web data used in the analysis

The final data set included 51 freshwater food webs, averaging 59 nodes and 196 links, sampled in 12 countries and six continents. See main text for information on how food webs were acquired and selected for this study.

**Table S1:** Information on the food webs filtered and used for SEM analysis

| Food Web | Country | Continent | Ecosystem | Type | Method | Reference |
|---|---|---|---|---|---|---|
| Afon Hafren 2005 | United Kingdom | Europe | Stream | Lotic | Gut-isotope | Layer et al. (2010) |
| Akatore A | New zealand | Oceania | River | Lotic | Gut-content | Thompson_Townsend (2004) |
| Allt a Mharcaidh | United Kingdom | Europe | Stream | Lotic | Gut-isotope | Layer et al. (2010) |
| Baoan Lake | China | Asia | Lake | Lentic | Modeling | Guo (2013) |
| Berwick | New zealand | Oceania | Stream | Lotic | Gut-content | Thompson_Townsend (2003) |
| Blackrock Stream | New zealand | Oceania | Stream | Lotic | Gut-content | Townsend et al. (1998) |
| Broad Stream | New zealand | Oceania | Stream | Lotic | Gut-content | Townsend et al. (1998) |
| Broadstone Stream | United Kingdom | Europe | Stream | Lotic | Gut-isotope | Layer et al. (2010) |
| Caballeros | Spain | Europe | Lake | Lentic | Gut-content | Sanchez (2015) |
| Canton Creek | New zealand | Oceania | Stream | Lotic | Gut-content | Townsend et al. (1998) |
| Catlins Stream | New zealand | Oceania | Stream | Lotic | Gut-content | Thompson_Townsend (2004) |
| Cimera | Spain | Europe | Lake | Lentic | Gut-content | Sanchez (2015) |
| Corrente River | Brazil | South America | River | Lotic | Modeling | Angelini et al. (2010) |
| Coweeta1 | United states | North america | Stream | Lotic | Gut-content | Thompson_Townsend (2003) |
| Coweeta17 | United states | North america | Stream | Lotic | Gut-content | Thompson_Townsend (2003) |
| Dargall Lane | United Kingdom | Europe | Stream | Lotic | Gut-isotope | Layer et al. (2010) |
| Dempsters Stream | New zealand | Oceania | Stream | Lotic | Gut-content | Townsend et al. (1998) |
| Duddon Pike Beck | United Kingdom | Europe | Stream | Lotic | Gut-isotope | Layer et al. (2010) |
| German Creek | New zealand | Oceania | Stream | Lotic | Gut-content | Townsend et al. (1998) |
| Grande De Gredos | Spain | Europe | Lake | Lentic | Gut-content | Sanchez (2015) |

| | | | | | | |
|---|---|---|---|---|---|---|
| Hardknott Gill | United Kingdom | Europe | Stream | Lotic | Gut-isotope | Layer et al. (2010) |
| Healy Creek | New zealand | Oceania | Stream | Lotic | Gut-content | Townsend et al. (1998) |
| Hongze Lake | China | Asia | Lake | Lentic | Modeling | Guo (2018) |
| Iceland Stream Is7 Aug 2008 | Iceland | Europe | Stream | Lotic | Gut-content | O'Gorman et al. (2012) |
| Iceland Stream Is8 Aug 2008 | Iceland | Europe | Stream | Lotic | Gut-content | O'Gorman et al. (2012) |
| Kye Burn | New zealand | Oceania | Stream | Lotic | Gut-content | Townsend et al. (1998) |
| Lake Biandantang | China | Asia | Lake | Lentic | Gut-content | Liu (2006) |
| Lake Hayq | Ethiopia | Africa | Lake | Lentic | Modeling | Fetahi et al. (2011) |
| Lake Huron | Canada | North America | Lake | Lentic | Modeling | Langseth (2014) |
| Lake Malawi | Tanzania | Africa | Lake | Lentic | Modeling | Nsiku (1999) |
| Lake Ontario | Canada | North America | Lake | Lentic | Modeling | Stewart & Sprules (2011) |
| Lake Shangshe | China | Asia | Lake | Lentic | Modeling | Li (2018) |
| Lake Taihu | China | Asia | Lake | Lentic | Modeling | Xu (2016) |
| Little Kye Burn | New zealand | Oceania | Stream | Lotic | Gut-content | Townsend et al. (1998) |
| Martins | United states | North america | Stream | Lotic | Gut-content | Thompson_Townsend (2003) |
| Mill Stream | United Kingdom | Europe | Stream | Lotic | Gut-isotope | Layer et al. (2010) |
| Mosendale Beck | United Kingdom | Europe | Stream | Lotic | Gut-isotope | Layer et al. (2010) |
| Mulgrave River | Australia | Oceania | River | Lotic | Gut-isotope | Rayner et al. (2010) |
| Northcol | New zealand | Oceania | Stream | Lotic | Gut-content | Thompson_Townsend (2003) |
| Old Lodge | United Kingdom | Europe | Stream | Lotic | Gut-isotope | Layer et al. (2010) |
| Paraguay River Bm Abaixo | Brazil | South America | River | Lotic | Modeling | Angelini et al. (2013) |
| Paraguay River Bm Acima | Brazil | South America | River | Lotic | Modeling | Angelini et al. (2013) |
| Parana River | Brazil | South America | River | Lotic | Modeling | Angelini & Agostinho (2005) |
| Peri Lake | Brazil | South America | Lake | Lentic | Gut-content | Peralta (2016) |
| Potreirinho Creek | Brazil | South America | Stream | Lotic | Gut-content | Motta & Uieda (2005) |
| Powder | New zealand | Oceania | Stream | Lotic | Gut-content | Thompson_Townsend (2003) |
| Stony Stream | New zealand | Oceania | Stream | Lotic | Gut-content | Townsend et al. (1998) |
| Sutton Stream | New zealand | Oceania | Stream | Lotic | Gut-content | Townsend et al. (1998) |
| Tonle Sap Lake | Cambodia | Asia | Lake | Lentic | Modeling | Chea (2016) |
| Troy | United states | North america | Stream | Lotic | Gut-content | Thompson_Townsend (2003) |
| Venlaw | New zealand | Oceania | Stream | Lotic | Gut-content | Thompson_Townsend (2003) |

**Web of science searched keyword for lake food webs:**

TI = (lake OR lakes OR pond OR ponds) AND TI = (food web OR food webs OR foodweb OR foodwebs OR food-web OR food-webs OR network OR networks) AND TI = (ecopath OR ecosim OR connectance OR topology OR structure OR architecture).

### 3. Land use intensity data for large waterbodies

For large water bodies, such as lakes and reservoirs, we obtained land use intensity values by considering the pixels within a 1 km buffer around the water area (Fig. S2).

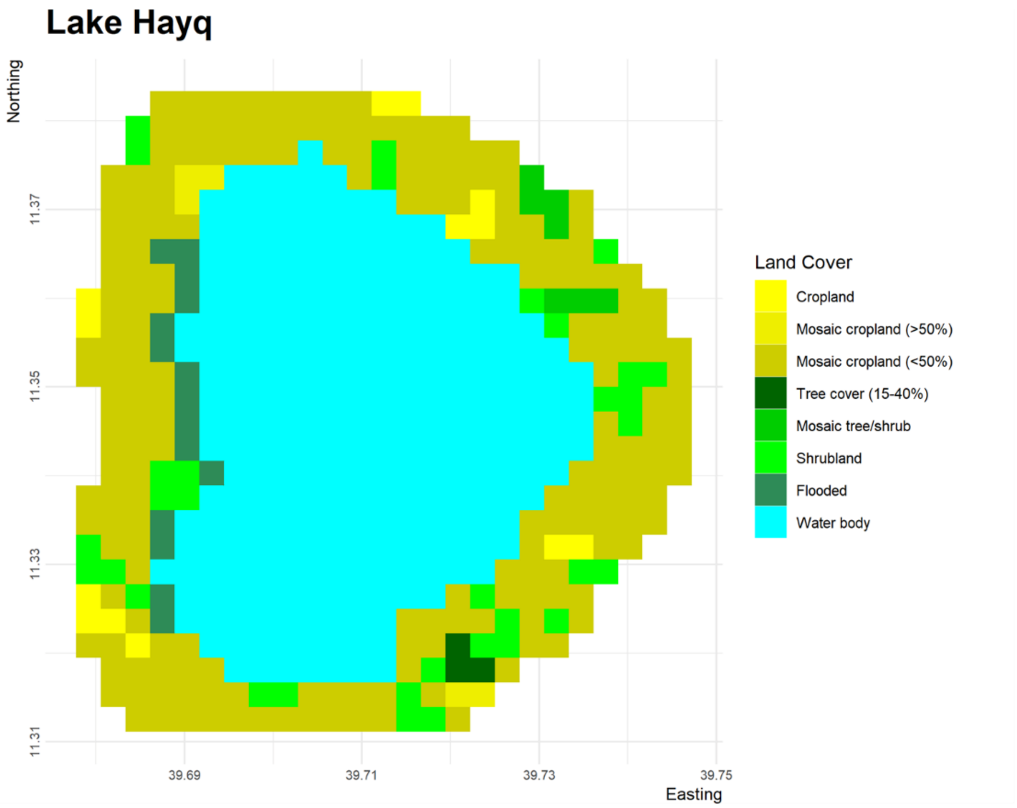


**Fig. S2**: Example of how land use intensity was obtained for large waterbodies (see main text for a description of calculation methods, land use classes and attributed pixel values).

**4. Sensitivity analysis:**

To ensure that differences in the number of lentic and lotic food webs compiled would not affect our model, we performed a sensitivity analysis in which we used the same number of lentic and lotic food webs by fitting the model on 999 networks randomly selected from our dataset.

.

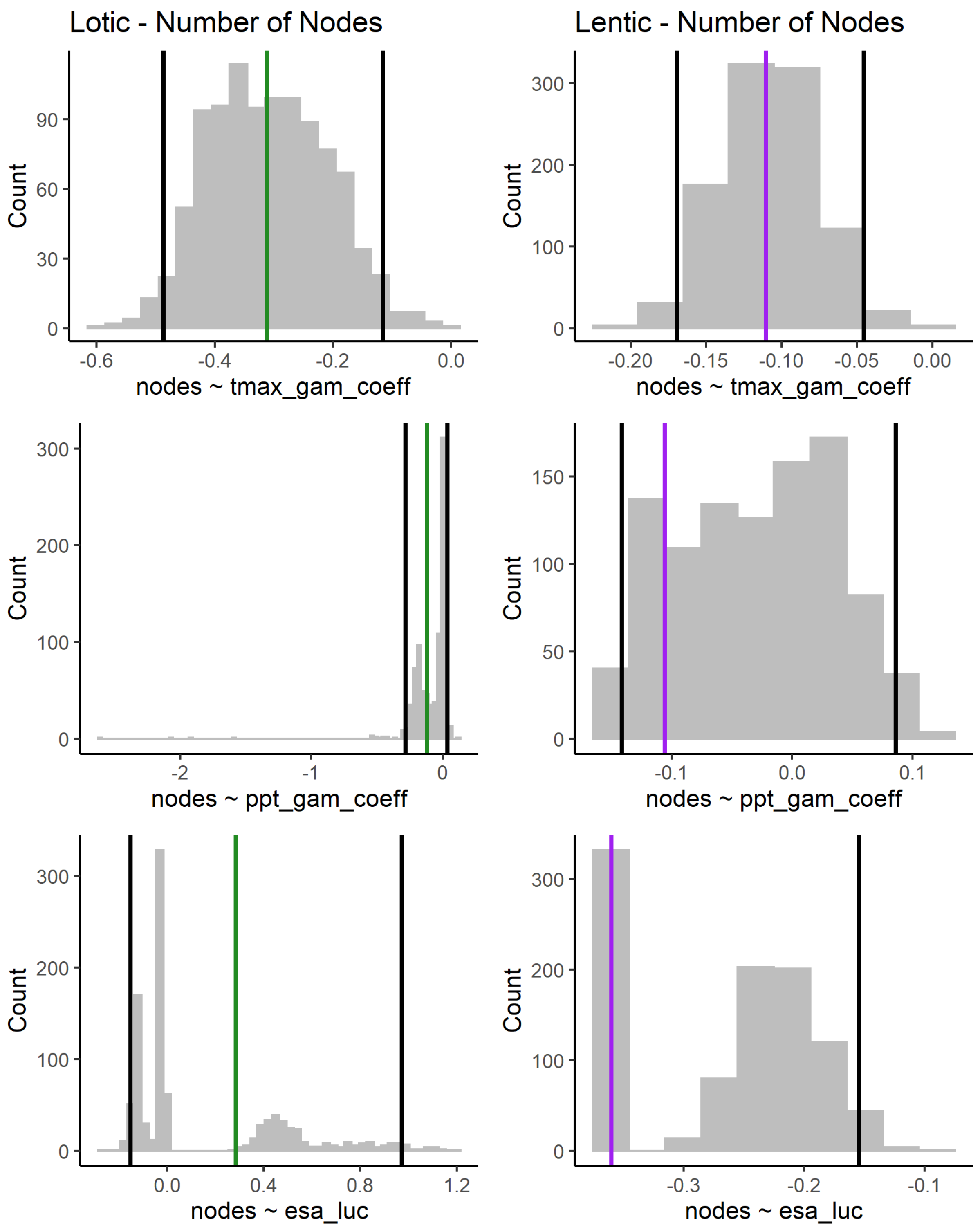

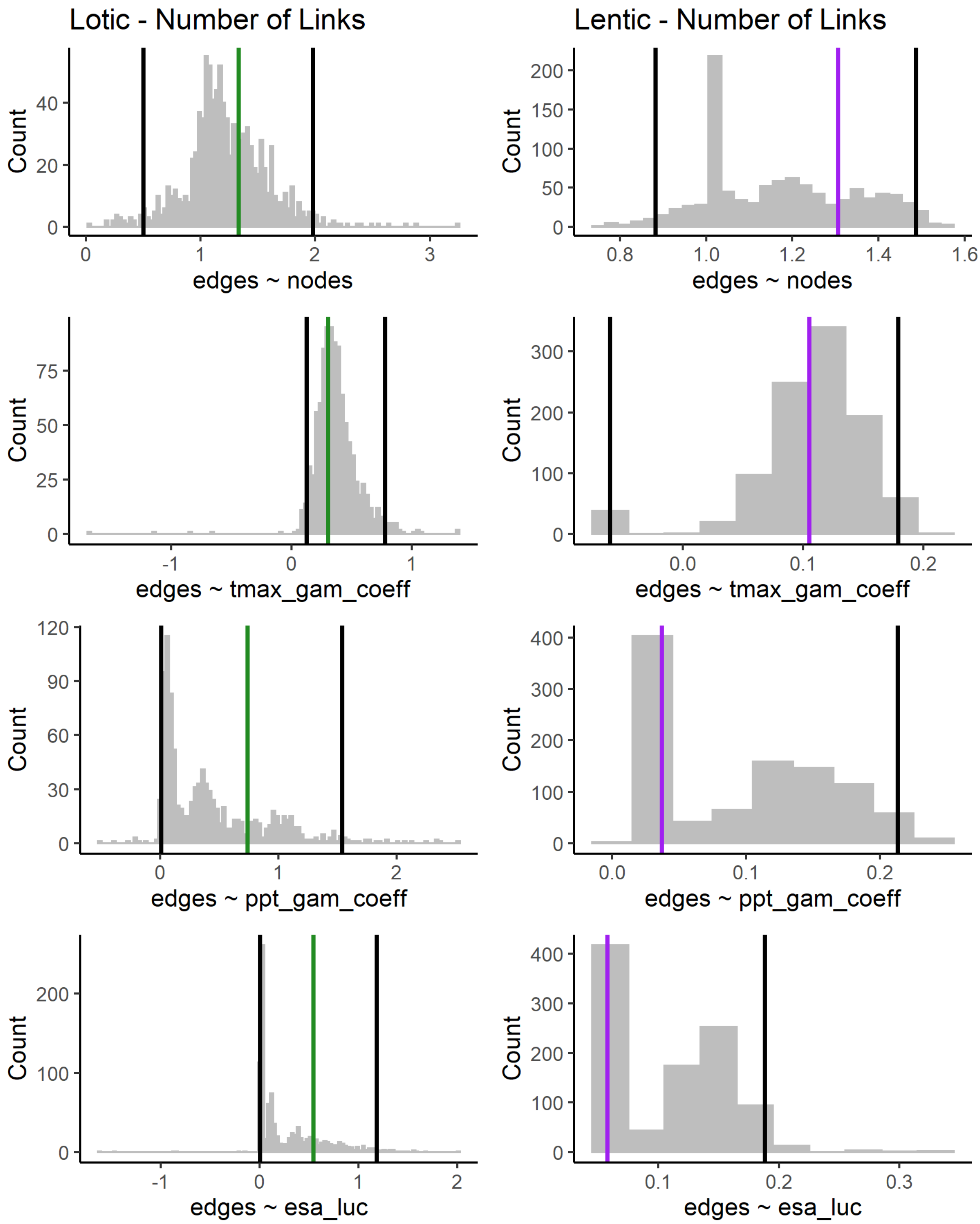

Lotic - Number of Links
Lentic - Number of Links
Count
edges ~ nodes
edges ~ tmax_gam_coeff
edges ~ ppt_gam_coeff
edges ~ esa_luc

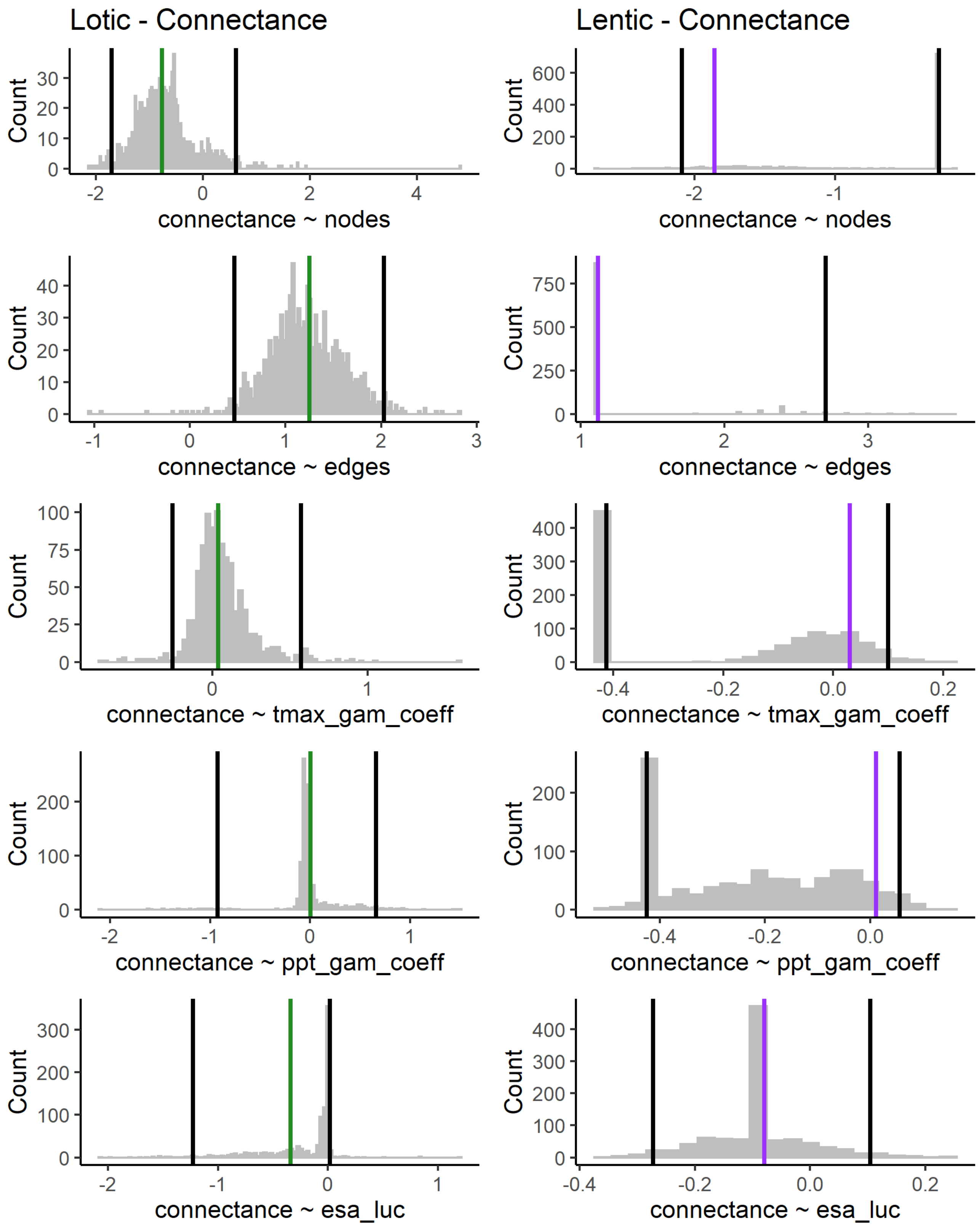

Lotic - Connectance
Lentic - Connectance
Count
connectance ~ nodes
connectance ~ edges
connectance ~ tmax_gam_coeff
connectance ~ ppt_gam_coeff
connectance ~ esa_luc

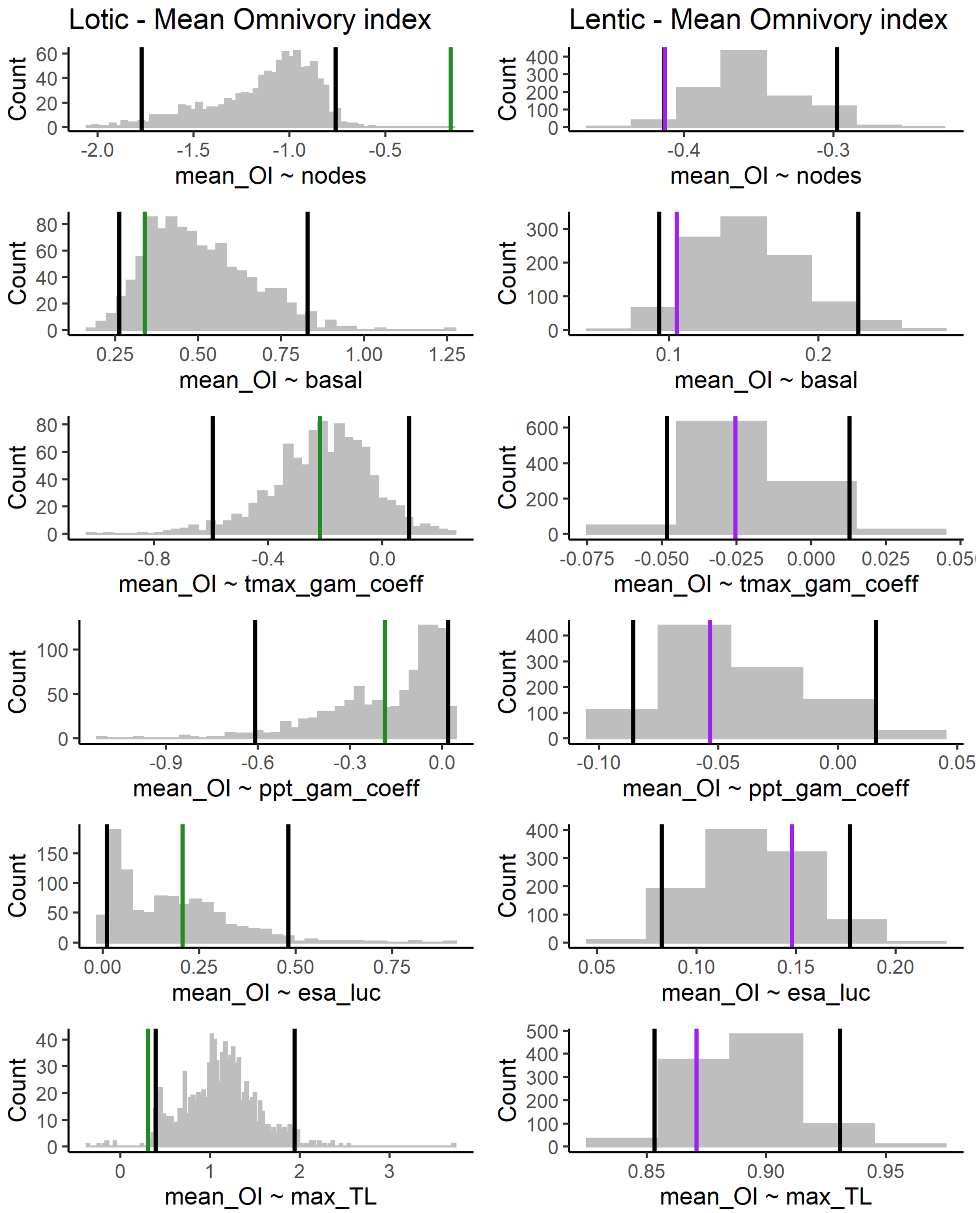

Lotic - Mean Omnivory index
Lentic - Mean Omnivory index
Count
mean_OI ~ nodes
mean_OI ~ basal
mean_OI ~ tmax_gam_coeff
mean_OI ~ ppt_gam_coeff
mean_OI ~ esa_luc
mean_OI ~ max_TL

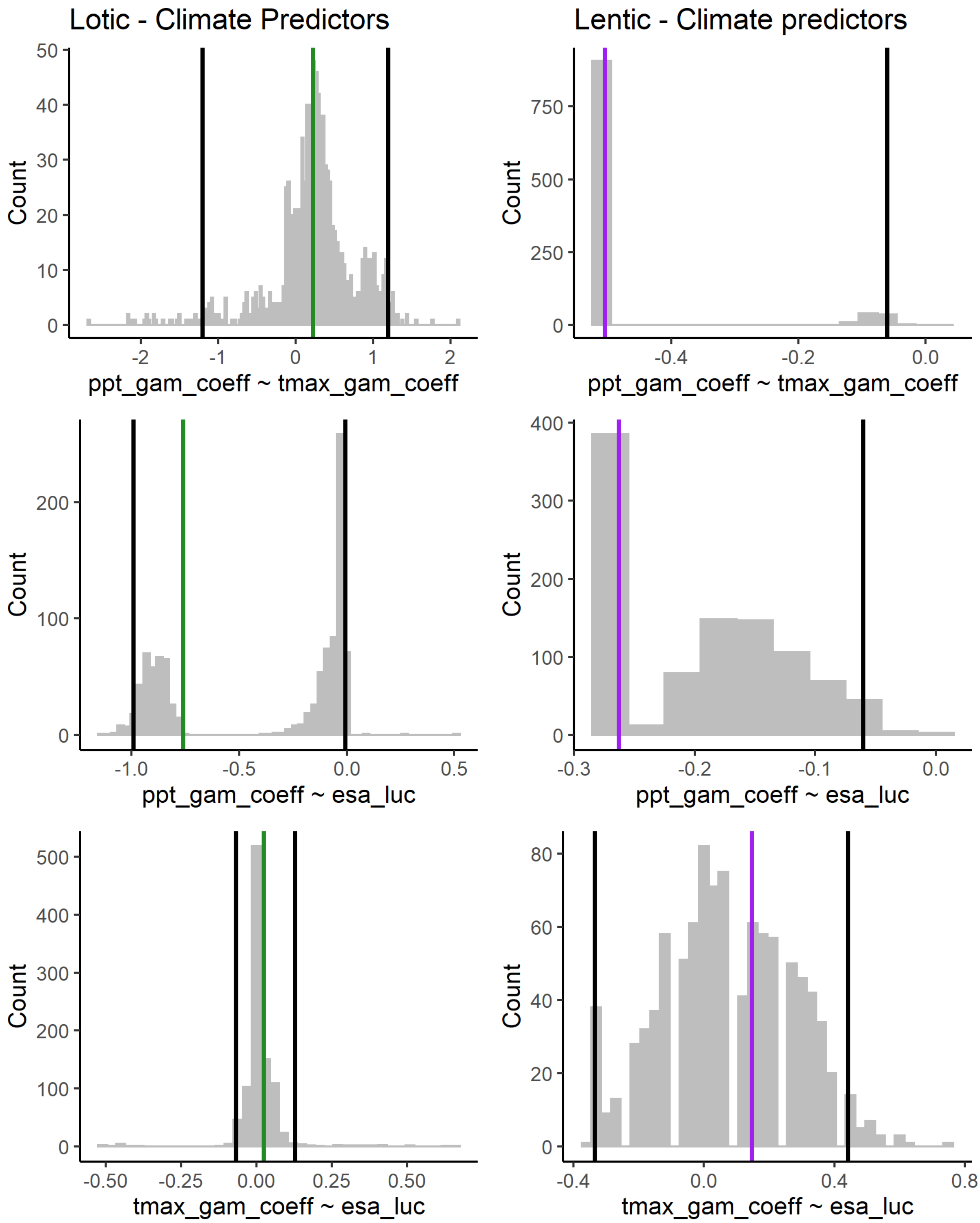


**Fig. S3**: Results of the sensitivity analysis performed with 999 lentic and lotic networks randomly selected from our dataset. Each graph refers to the relationship between exogenous and endogenous variables in our model.
